# Relevance of coral geometry in the outcomes of the coral-algal benthic war

**DOI:** 10.1101/327031

**Authors:** Emma E. George, James Mullinix, Fanwei Meng, Barbara Bailey, Clinton Edwards, Ben Felts, Andreas Haas, Aaron C. Hartmann, Benjamin Mueller, Jim Nulton, Ty N.F. Roach, Peter Salamon, Cynthia B. Silveira, Mark J.A. Vermeij, Forest L. Rohwer, Antoni Luque

## Abstract

Corals have built reefs on the benthos for millennia, becoming an essential element in marine ecosystems. Climate change and human impact, however, are favoring the invasion of non-calcifying benthic algae and reducing coral coverage. Corals rely on energy derived from photosynthesis and heterotrophic feeding, which depends on their surface area, to defend their outer perimeter. But the relation between geometric properties of corals and the outcome of competitive coral-algal interactions is not well known. To address this, 50 coral colonies interacting with algae were sampled in the Caribbean island of Curaçao. 3D and 2D digital models of corals were reconstructed to measure their surface area, perimeter, and polyp sizes. A box counting algorithm was applied to calculate their fractal dimension. The perimeter and surface dimensions were statistically non-fractal, but differences in the mean surface fractal dimension captured relevant features in the structure of corals. The mean fractal dimension and surface area were negatively correlated with the percentage of losing perimeter and positively correlated with the percentage of winning perimeter. The combination of coral perimeter, mean surface fractal dimension, and coral species explained 19% of the variability of losing regions, while the surface area, perimeter, and perimeter-to-surface area ratio explained 27% of the variability of winning regions. Corals with surface fractal dimensions smaller than two and small perimeters displayed the highest percentage of losing perimeter, while corals with large surface areas and low perimeter-to-surface ratios displayed the largest percentage of winning perimeter. This study confirms the importance of fractal surface dimension, surface area, and perimeter of corals in coral-algal interactions. In combination with non-geometrical measurements such as microbial composition, this approach could facilitate environmental conservation and restoration efforts on coral reefs.

## INTRODUCTION

Corals use energy derived from photosynthesis and heterotrophic feeding to build reefs. This has enabled corals to dominate the battle for light and space on the reef benthos for millennia (Kaandorp & Kubler, 2001). However, the combination of overharvesting of herbivorous fish, increased nutrient runoff from land (eutrophication), and ocean warming is stimulating the growth of non-calcifying algae at the expense of corals world-wide (Alevizon & Porter, 2015). The increase in algal coverage is re-routing the energy to alternative trophic pathways that are enhancing the dominance of algae through positive feedback loops, for example, invigorating the growth of opportunistic and virulent microbes at the coral-algal interface (Kline et al., 2006; Smith et al., 2006; Dinsdale & Rohwer, 2011, Silveira et al. 2015). As algal density increases on reefs, competitive coral-algal interactions are becoming more frequent (Barott et al., 2012a,b; Dinsdale & Rohwer, 2011; Haas et al., 2011), and, in order to preserve and restore coral reefs, it is crucial to understand the key factors that determine the outcomes of these interactions.

While there has been significant study into the effects of nitrification and changes in herbivore biomass on coral-algal interactions, results have been somewhat equivocal (Smith et al. 2001, McCook et al. 2001, Burkepile et al. 2006, Rasher et al. 2012). This suggests that other factors such as coral colony conditions may contribute to the outcome of coral-algal interactions. In fact, according to the DDAM (DOC-Disease-Algae-Microbes) hypothesis, dissolved organic carbon (DOC) released by fleshy algae stimulates the growth of opportunistic microbes at the coral-algal interface (Dinsdale & Rohwer, 2011). In combination with a shift to inefficient microbial metabolic pathways, this is suggested to lead to hypoxic conditions at the coral-algal interface, weakening and killing coral tissues (Haas et al., 2013; Roach et al., 2017). Simultaneously, the outcome of a competitive interaction with benthic algae depends on the relative algae overgrowth rate as well as the percentage of the coral perimeter in contact with macroalgae (Lirman, 2001). Thus, the coral perimeter and the ability to defend it must be a key factor in determining the coral-algal interaction outcome.

A coral colony consists of multiple clonal polyps that are connected by the coenosarc tissue. Polyps along the perimeter of the colony interact with invading non-calcifying algae as well as other benthic organisms (Jackson, 1977 & 1979; Buss & Jackson, 1979; Meesters, Wesseling & Bak, 1996). At any competitive interaction zone a coral can either overgrow (win), be overgrown (lose), or neither overgrow nor be overgrown (neutral) by the interacting species (Figure S1A) (Jackson & Winston, 1982; Barott et al., 2012b; Swierts & Vermeij, 2016).

Defending the perimeter requires the allocation of resources. The energy obtained from photosynthesis—carried out by endosymbiotic algae—and heterotrophic feeding (Porter, 1976) is then distributed throughout the colony using the coenosarc tissue (Rinkevich & Loya, 1989; Oren et al., 1997; Henry & Hart, 2005; Schweinsberg et al., 2015). As the colony’s surface area increases so does its potential for nutrient acquisition and distribution (Oren et al., 2001). Thus, coral surface area should be another key factor in determining the coral-algal interaction outcome.

The resource availability hypothesis (RAH) (Endara & Coley, 2011) predicts that fast growing corals will rely on clonal growth strategies to indirectly outcompete the invading algae. This explains the resilience observed among branching corals, which invest the resources acquired from their large surface areas to grow new polyps rather than to protect their small perimeters (Swierts & Vermeij, 2016). In contrast, RAH predicts that slow growing species tend to face more encounters with competitors and will invest more resources in protecting their perimeters. This has been confirmed for slow growing corals like encrusting and massive corals (Swierts & Vermeij, 2016).

The morphology and size of these slow growing corals have been linked to corals’ natural competitive edge against most algal groups (Porter, 1976; Tanner, 1995). Massive corals have relatively lower perimeter-to-surface area ratios and demonstrate greater resilience to algal overgrowth compared to encrusting corals with large perimeter-to-surface area ratios (Hughes 1989; Tanner 1995; Lirman 2001). A coral-algal survey in the Line Islands observed that small and large corals were more effective winning against algae than medium sized corals (40–80 cm) (Barott et al. 2012b). In contrast, in the South China Sea it was observed that medium size corals won more often than small and large corals (Swierts & Vermeij, 2016). Thus, the influence of the geometrical properties in the outcome of the coral-algal interaction remains unclear.

The accurate measurement of the perimeter and surface area in natural objects, however, is usually challenged by the presence of *fractality* (Mandelbrot, 1967, 1977, 1983). Fractals are non-smooth objects that display similar patterns across multiple scales. This makes the perimeter length and surface area to depend on the resolution of the measurement. In particular, the values follow a power law of the scale with an exponent related to the perimeter and surface’s fractal dimensions, respectively, (Falconer 2003, Okie, 2013) (see Eq. (S1) in Methods). Higher fractal dimensions lead to more convoluted surfaces or perimeters with a larger number of wrinkles and textures that increase the effective surface and perimeter of corals (Falconer, 2003; Okie, 2013). Previous studies found fractality among corals at different scales (Basillais, 1997; Bradbury & Reichelt, 1983; Knudby & LeDrew, 2007; Martin-Garin et al., 2007; Mark, 1984; Purkis et al., 2006; Reichert et al., 2017; Zawada & Brock, 2009), but the measurements at the coral colony scale of interest in the present study were inconclusive (Mark 1984).

Here we hypothesize that larger fractal dimensions and smaller perimeter-to-surface area ratios would favor corals when facing competitive interactions with algae. To characterize the fractal dimension accurately, high-resolution images of corals were necessary (Young et al., 2017), so we applied new imaging and computer rendering technologies to obtain a systematic and accurate analysis of coral geometry in the 1 mm to 1 m range.

## METHODS

### Field sampling

Photographs of 50 coral colonies in the Caribbean island of Curaçao were taken by SCUBA diving using a Canon Rebel T4i with a 35-mm lens and two Keldan 800 lumen video lights to illuminate the corals uniformly. An in-reef ruler was photographed to set the scale for the digital models; the ruler was placed along the interface of the coral colonies. Additionally, the perimeters of five coral colonies were measured in the field using a chain-link method, using links of sizes 1.5 cm, 5.5 cm, and 10.5 cm. See Supplementary Material for additional details.

### 2D perimeter models and competition outcomes

High-resolution, overlapping images of coral perimeters were stitched together to build a 2D perimeter model (see Figures 1 and 2) using Globalmatch and Guimosrenderer software (Gracias & Santos-Victor, 2000, 2001; Lirman et al., 2007, 2010). The minimum threshold resolution was ∼0.5 mm. The interaction zones were outlined in separate RGB channels using Adobe^®^ Photoshop^®^ *CC* 2014 (Figures 1 and 2): red (coral losing), green (coral winning), and blue (neutral). The fraction of red, green, and blue pixels was used, respectively, to obtain the percentage of losing (%L), winning (%W), and neutral (%N) interactions around a coral perimeter. See Supplementary Material for additional details.

**Figure 1.**
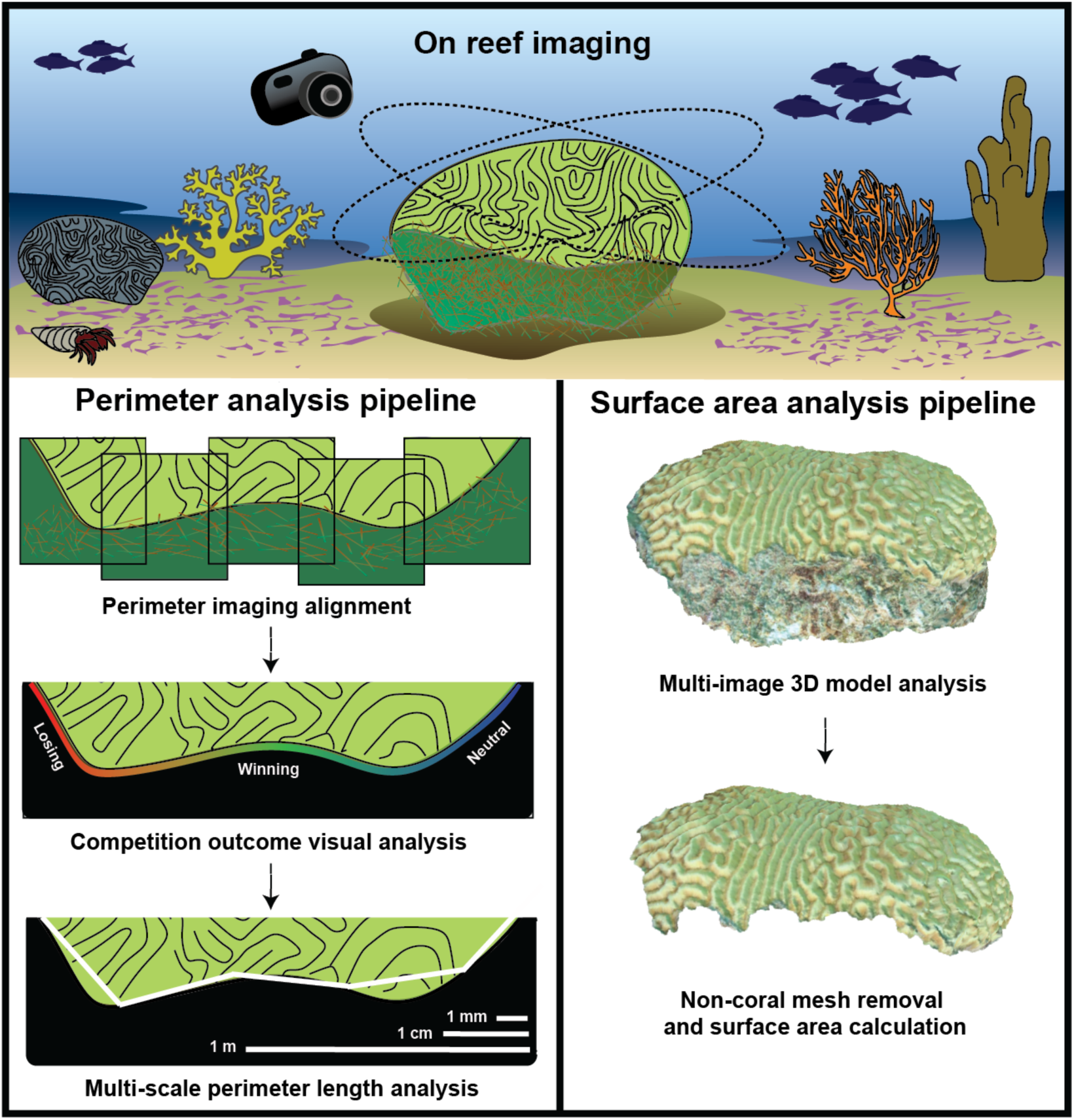
Coral geometry methods. (Top panel) Corals were photographed from different angles and distances. (Bottom left panel) Close range pictures were stitched together to generate a high-resolution 2D perimeter model. The interactions along the coral perimeter were outlined and the perimeter lengths were measured over a 0.1 mm to 1 m scale range. (Bottom right panel) Farther range pictures were processed to create the 3D coral models. Models were calibrated with an in-reef reference; non-coral mesh was removed to measure the coral surface area.

**Figure 2.**
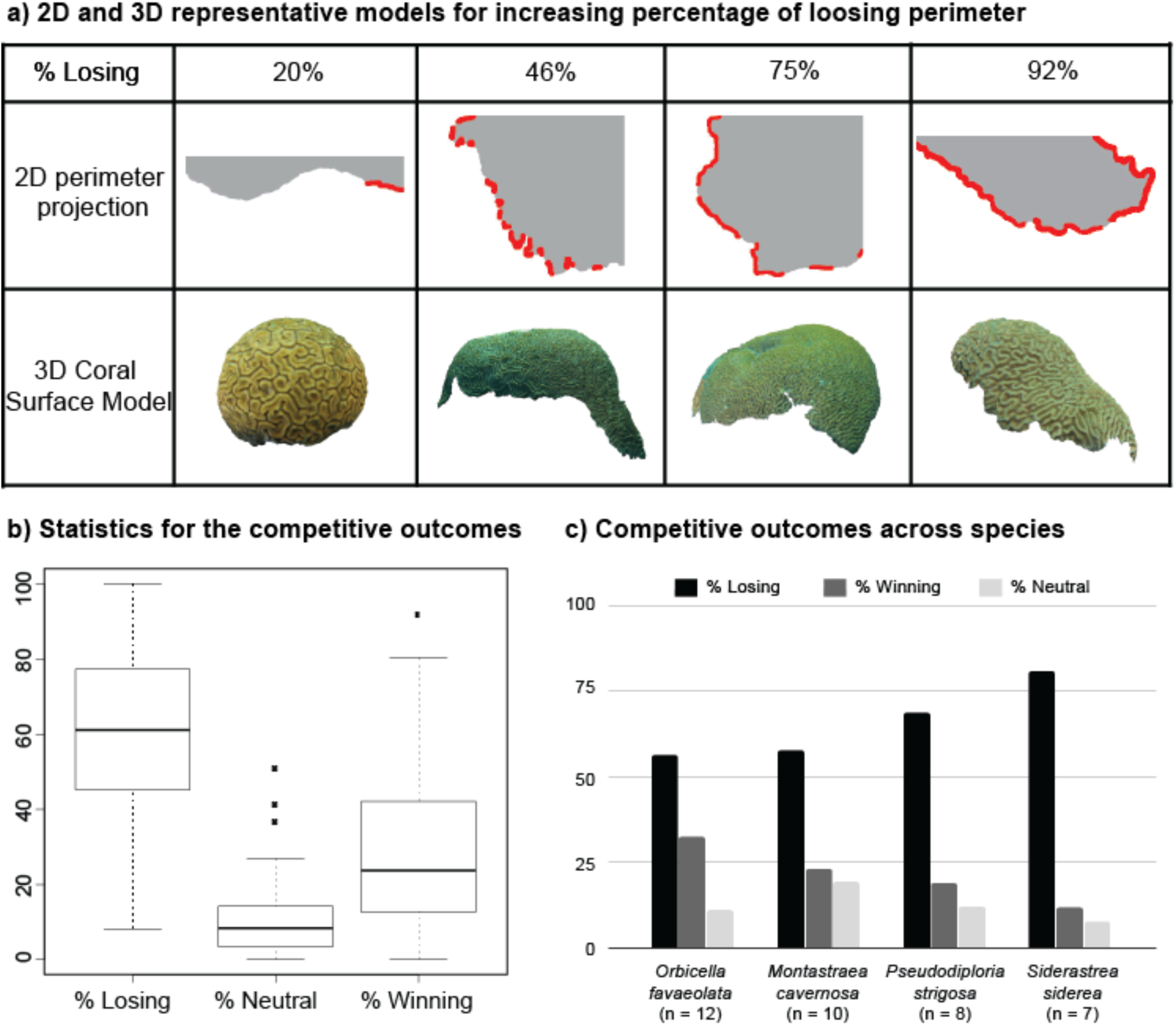
Coral models and statistics for competitive outcomes. a) 2D and 3D coral models for different percentages of losing perimeter (%L). The 2D models highlight the losing regions in red. b) Box plot for the three perimeter outcomes: losing (%L), neutral (%N), and winning (%W). The middle line corresponds to the median, the range of the box contains from the 25^th^ to the 75^th^ percentile, and each whisker is the minimum (in absolute value) between the 150% interquartile range (IQR) and the value of the most extreme point in that side of median. Outliers exceeding the whiskers are included (Table S5). b) Competitive outcomes for species that were sampled in five or more colonies. The bars correspond to the average percentage of losing perimeter (black), average percentage of winning perimeter (dark grey), and average percentage of neutral perimeter (light grey).

### 3D coral models

Autodesk^®^ ReMake^®^, 2016 was used to create 3D coral models (Burns et al., 2015; Leon et al., 2015) (Figures 1 and 2) to facilitated the accurate measurement of geometric properties of corals, e.g., perimeter, surface area, and volume at multiple scales (Naumann et al., 2009 & Lavy et al., 2015). The resolution of the models ranged from 1.6 mm to 49 mm with an average of 11 mm. See Supplementary Material for additional details.

### Perimeter and surface fractal dimensions

The fractal dimension was calculated using a box counting method (Falconer 2003). The logarithm of the number of boxes was plotted against the logarithm of the box size, and the fractal dimension *D* was extracted from the slope of the linear regression using Eq. (S1) (Figure S2). The 95% confidence intervals for the fractal dimension was calculated using a Monte-Carlo non-parametric bootstrap resampling method. The method was tested for the following fractal objects: the Koch curve, Sierpinski triangle, Menger sponge, and kidney vasculature. This lead to an error on the 1–3% range using five bisections (Table S1). The upper value of this range was used as the theoretical error for the fractal dimension. The perimeter fractal dimension (DP) was calculated from the 2D high-resolution models, which allowed a minimum of ten bisections in the algorithm. The surface fractal dimension (DS) was calculated from the 3D high-resolution models, which allowed a minimum of five bisections in the algorithm. The null hypotheses DP ≠ 1 and DS ≠ 2 were evaluated using the nonparametric sign test. See Supplementary Material for additional details.

### Coral geometric properties: perimeter, surface area, volume, and polyp size

The perimeter, surface area, and volume of the 3D models were calculated with the mesh report tool in Autodesk^®^ Remake^®^, 2016. This approach was previously tested in Naumann et al., 2009 and Lavy et al., 2015. The perimeter of the high-resolution 2D models was obtained using a Richardson algorithm. The values were compared with field values using three physical chain-links (1.5 cm, 5.5 cm, and 10.5 cm) with an errors of 14.5%, 17.5%, and 19.7%, respectively (Table S2). This discrepancy was reasonable taking into the account the projection on the model and the measurement field error. The 2D perimeter used in the analysis was obtained using a 1 mm ruler in the Richardson algorithm. Polyp diameters were also measured from the 2D models using ImageJ 1.47v and averaging 10 polyp diameters per colony. See Supplementary Material for additional details.

### Correlation with single variables

A linear regression (least squares method) was used to compare the percentages of losing (%L) and winning (%W) perimeter with respect depth (d), polyp diameter (Pd), volume (V), surface area (SA), surface area-to-polyp area ratio (SApolyp), perimeter fractal dimension (DP), surface fractal dimension (DS), 2D perimeter obtained from Richardson’s algorithm (PR), 3D perimeter length (P3D), 3D perimeter-to-polyp size ratio (Ppolyp), 2D perimeter-to-surface area ratio (PR/SA), and 3D perimeter-to-surface area ratio (P3D/SA) (Table S6). The neutral interactions were a small fraction and were not studied in detail.

### Statistical learning: Random forest

The package randomForest (Liaw & Wiener, 2002) was used to analyze the response of percentage of perimeter losing (%L) and winning (%W) as function of the 13 variables listed above. The package rfPermute (Archer 2016) was used to estimate the significance of importance and p-value. metrics by permuting the dependent variable, producing a null distribution of importance metrics, and calculating the p-value for each predictor variable. The initial global analysis included 13 input variables: Species, depth, polyp diameter, volume, surface area, volume to surface area ratio, surface to polyp diameter square ratio, projected perimeter length, 3D perimeter length, 3D perimeter length to polyp diameter ratio, projected perimeter to surface area ratio, 3D perimeter to surface area ratio, perimeter fractal dimension, and surface fractal dimension. Both rfPermute and randomForest were run five times and averaged separately to rank the variables independently based on the mean increase accuracy error %IncMSE values. The analysis combining the top ranked variables in groups of three, and the combination leading to the largest variance explained was selected for further analysis. The hierarchical visualization of these variables was obtained using the rpart package in R (Terry 2017) for %L and %W. See Supplementary Material for additional details.

### Coral geometric properties across Curaçao regions

The corals sampled (n) were grouped in three geographical regions in the island of Curaçao: East (n=9), Central (n=37), and West (n=4)—see Figure S3. The four main geometrical indicators for the percentage of losing and winning interactions (fractal surface dimension, surface area, perimeter length, and perimeter-to-surface area ratio) were compared using boxplots.

## RESULTS

### Coral-algal competition outcomes

On average, coral displayed 60% losing, 29% winning, and 11% neutral interactions along the perimeter with algae (Figures 2a and 2b, and Table S5). Among species that were sampled in five or more colonies, *S. siderea* displayed the largest percentage of losing perimeter (81%), followed by *P. strigosa* (69%), *M. cavernosa* (58%), and *O. faveolata* (56%) (Figure 2c). The species followed the inverse trend regarding the percentage of winning perimeter: *O. faveolata* (33%), *M. cavernosa* (23%), *P. strigosa* (19%), *S. siderea* (12%). The percentage of neutral perimeter was smaller and followed a different trend: *M. cavernosa* (19%), *P. strigosa* (12%), *O. faveolata* (11%), *S. siderea* (8%). Thus, corals were generally losing, and the neutral regions represented the smallest fraction of the competitive outcomes. On average, *S. siderea* was the most vulnerable species, while *O. faveolata* was the most resilient.

### Coral perimeter and surface fractal dimension

The perimeter fractal dimension, DP, for the 50 corals was very close to the Euclidean value, D = 1, and it was contained within the 5% to 95% confidence interval for all corals but three (CUR34, CUR54, and CAS142) (Figure 3a). When considering the theoretical error of the box counting method (3%), these three cases were compatible with the Euclidean value: CUR34 (DP = 1.00 ± 0.03), CUR54 (DP = 0.99 ± 0.03), and CAS142 (DP = 0.99 ± 0.03). The average fractal dimension was <DP> = 0.999± 0.03 (SE), and the nonparametric sign test evaluated if the individual perimeters were non-fractal as a whole (null hypothesis, *D* ≠ 1), yielding a p-value of 0.013. This was a conservative analysis, and incorporating the theoretical error (3%) would reduce this p-value even further. Thus, the dimensions of coral perimeters were non-fractal. The perimeters were also analyzed visually, when comparing high (DP = 1.01±0.03), medium (DP = 0.999±0.005), and low (DP = 0.988±0.008) fractal dimensions (±SE), no salient geometric feature distinguished them (Figure 3b).

**Figure 3.**
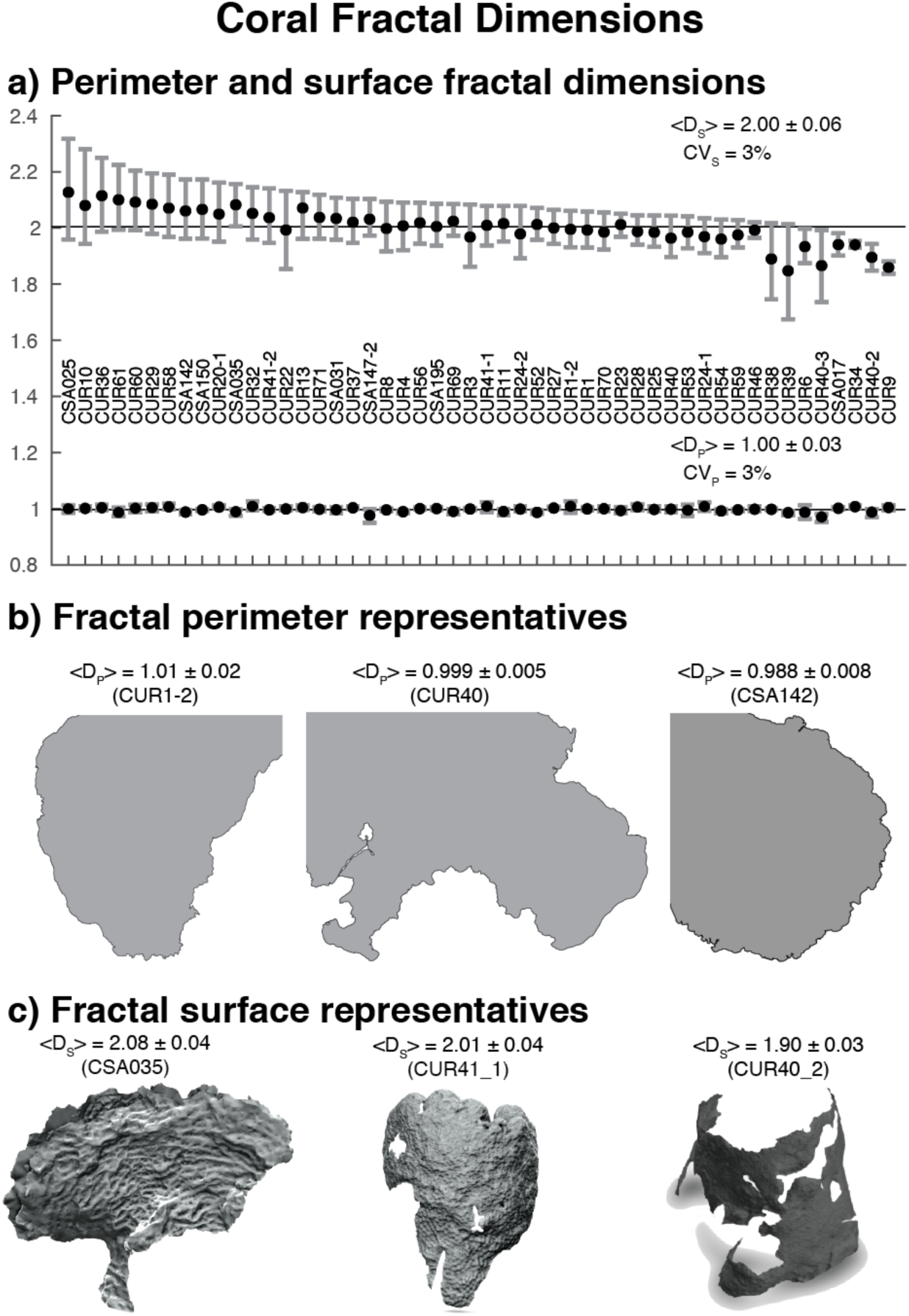
Coral fractal dimensions. a) Surface fractal dimensions (top) and perimeter fractal dimensions (bottom) for all specimens reconstructed digitally. The plot includes the mean (black dot), 5 to 95% confidence intervals (whiskers), and the label associated to each coral. A solid line provides a reference for the topological dimensions: D = 1 (perimeter) and D=2 (surface). The plot includes also the mean values for the fractal dimension of the perimeter (<D_P_>) and the surface (<D_S_>) (± standard deviation) and their respective coefficients of variation (CV = standard deviation / mean * 100). Panels b) and c) display coral representatives associated with high, medium, and low fractal dimension for the perimeter (b) and the surface (c).

The surface areas and surface fractal dimensions were measured for 50 corals within the 1 mm to 1 m range using the 3D coral models (Figure 3a). The 5% to 95% confidence intervals included the Euclidean surface dimension, D=2, for all corals except four: CSA017 (CI: 1.94– 1.98), CUR34 (CI:1.94–1.95), CUR40-2 (CI:1.90–1.94), CUR9 (CI:1.84–1.88). When considering the theoretical error of the box counting algorithm (∼3%), CSA017 (DS = 1.94 ± 0.06) and CUR34 (DS = 1.94 ± 0.06) were compatible with the Euclidean value, while CUR40-2 (DS = 1.90 ± 0.06) and CUR9 (DS = 1.86 ± 0.06) remained slightly lower. The average fractal dimension was <D_s_> = 2.00 ± 0.06 (±SE) The nonparametric sign test evaluated if the coral surfaces were fractal (null hypothesis, *D* ≠ 2); this yielded a p-value of 1.212e-7*** using the statistical confidence interval. This p-value would have been even smaller if the theoretical error was included. Thus, overall coral surfaces were statistically non-fractal. Figure 3c compares corals with high (2.08 ± 0.04), medium (2.01 ± 0.04), and low (1.90 ± 0.03) mean surface fractal dimensions. Corals with high surface fractal dimension had no holes and their surface texture was more rugose; corals with low fractal dimension instead displayed holes, peninsulas, and smoother surfaces. Thus, coral surfaces were statistically non-fractal, but the mean fractal value captured distinguishable geometrical features.

### Relationship between outcomes and individual geometric variables

The absence of fractality in corals facilitated the measurement of the geometric properties at a single (high-resolution) scale. The percentage of losing perimeter (%L) was studied as a function of geometric and biological variables using linear regression analysis (see Figure S5 and Table S6). The percentage of losing perimeter (%L) was negatively correlated with the surface area (slope = 8.6±4.2 1/log10(cm2), R^2^ = 0.09, p-value=0.045*) and the surface fractal dimension (slope = –145±45, R^2^ = 0.18, p-value = 0.0021**). The opposite was observed for the percentage of winning perimeter (%W): surface area (slope = 8.6±4.2 1/log10(cm2), R^2^ = 0.08, p-value=0.045*) and surface fractal dimension (slope = 144±45, R^2^ = 0.18, p-value = 0.0023**). This is due to %W being negatively correlated with %L (slope = –0.9 ± 0.1, R^2^ = 0.8, p-value = 2.2×10^−16^***) (Figure S4). The percentage of neutral perimeter (%N) was discarded due to its low values (Figure 2b). Thus, two surface properties (area and fractal dimension) were directly correlated with the coral competition outcomes. The fractal dimension displayed the strongest correlation, but only captured 18 % of the variance (R^2^ = 0.18), and no variables related to the perimeter showed a direct correlation with the outcomes.

### Importance of combined geometric variables in coral-algal competition outcomes

The combined effect of coral geometric properties in predicting coral-algal competitive outcomes was analyzed using random forest, which estimated the average percentage increase of mean squared error <%IncMSE> in predicting the losing perimeter (%L) for each coral feature (see Figure S6a). The variance explained using all variables was 4.3 ± 0.6 % (SE). The surface fractal dimension was the most important predictor, and the only one selected statistically against the null hypothesis by rfPermute (p-value < 0.05). The following variables—listed with decreasing importance—were the 3D perimeter, surface area, and perimeter-to-surface ratio. The lowest ranked predictor was the mean perimeter fractal dimension.

The top ranked variables were then combined separately and analyzed again using the random forest statistical model (Table S6). The optimal combination was surface fractal dimension, 3D perimeter, and species. This explained 18.7 ± 0.5% (SE) of the variance, and the 3D perimeter (<%IncMSE> = 11.0 ± 0.3, p-value = 0.036* ± 0.009) and the surface fractal dimension (<%IncMSE> = 11.0 ± 0.4, p-value = 0.021* ± 0.007) were both equally important and statistically significant (p-value < 0.05) (Figure S6a). These two variables alone, however, explained only ∼8% of the variance. Combinations with other geometric variables, like the perimeter-to-surface ratio, led to ∼17% variance explained (see Table S6). Thus, coral geometry alone explained up 17% of the percentage of losing perimeter, and the surface fractal dimension and 3D perimeter were the most relevant variables.

An analogous analysis was done for the %Winning outcome. Figure S6b plots the input variables ranked as a function of their average percentage increase of mean squared error <%IncMSE>. Surface area, perimeter-to-surface area ratio, and 3D perimeter were the better-ranked variables, although only the surface area and 3D perimeter to surface area ratio had a significant p-value (0.05). The fractal surface dimension occupied a middle-ranked position, despite displaying a strong direct correlation with %W (Figure S5b); the perimeter fractal dimension was again the least relevant variable. The variance explained using all variables was 19.6% ± 0.9% (SE). As in the %Losing case, the most relevant variables were re-analyzed separately (Table S6). The optimal combination corresponded to the 3D perimeter to surface ratio, 3D perimeter, and surface area. This explained 26.6% ± 0.5% of the variance. The 3D perimeter to surface area ratio (12.0% ± 0.4%, p-value = 0.028* ± 0.007) and the 3D perimeter (10.6% ± 0.3%, p-value = 0.020* ± 0.004) were the most important and significant variables. The surface area had a similar value but the p-value was slightly larger (p-value = 0.059 ± 0.010). The geometrical properties of corals explained ∼25% of the variability of %Winning outcomes, and the perimeter to surface area ratio was the strongest predictor.

### Hierarchical analysis of coral outcomes and coral geometry

To gain insight on the relationship between coral geometrical properties and coral-algal competitive outcomes, regression tree models (rpart package in R, Terry 2017) were generated using the most relevant variables selected by random forest for %L and %W (see previous sections).

For the percent losing case (%L), the nodes of the regression tree corresponded to the surface fractal dimension and 3D perimeter (see Figure 4a). Corals with a fractal dimension DS < 2 had a higher %L and were classified on the left side of the tree. Among those, corals with 3D perimeters smaller than 318 cm formed the group with the largest percentage of losing perimeter, <%L> = 79%. For the group with DS > 2, a 3D perimeter larger than 549 cm led to the cluster with the lowest percentage of losing perimeter, <%L> = 44%. Figure 4a also displays the %L as a function of the 3D perimeter and surface fractal dimension. The sectors represent the regions selected by the tree. As expected, the bottom-left sector (small DS and small perimeter) had the highest value of percentage losing perimeter, while the top-right sector (large DS and perimeter) had the smallest percentage of losing perimeter.

**Figure 4.**
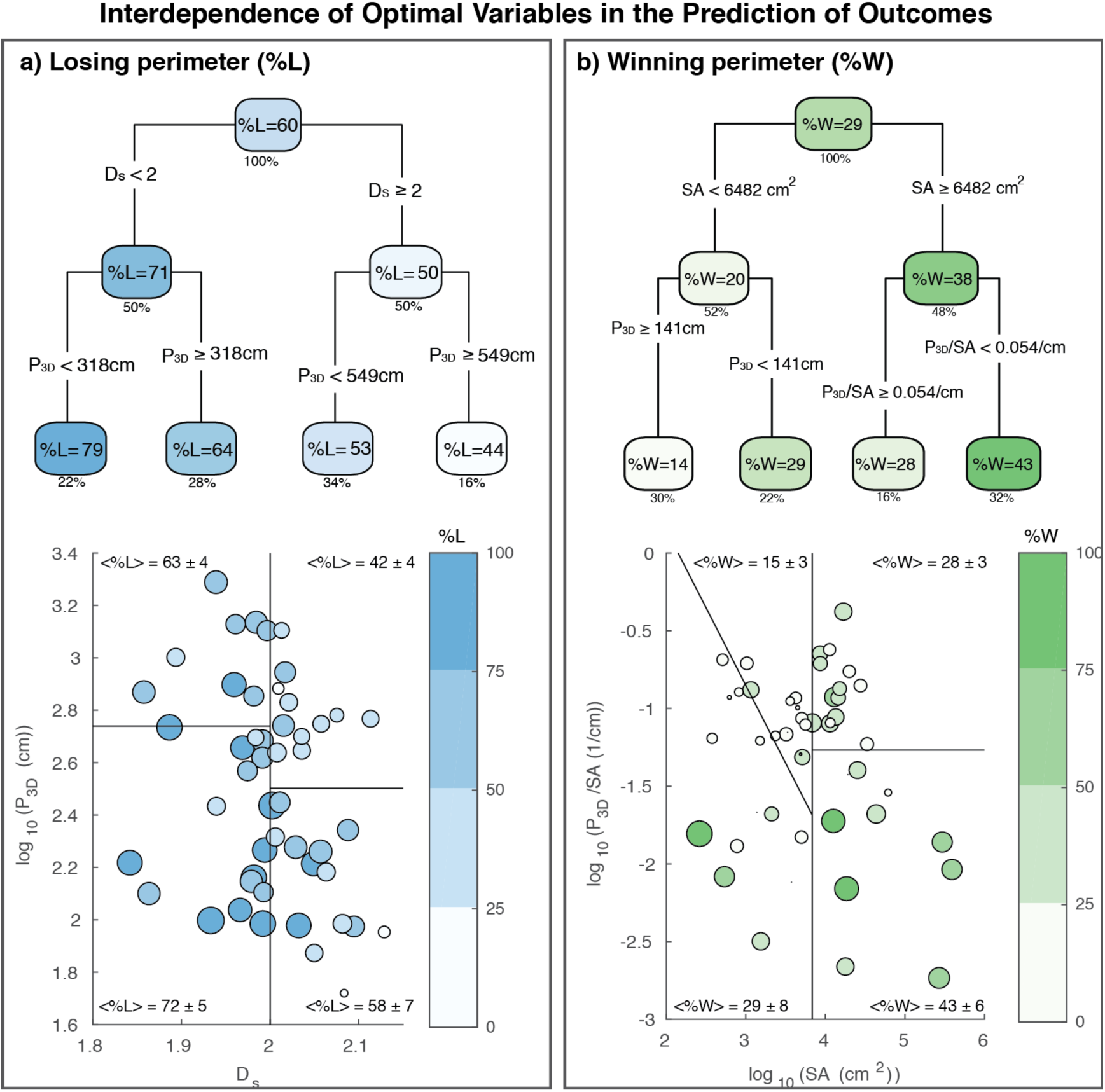
Interdependence of optimal variables in the predictions of outcomes. a) A regression tree (top panel) generated for the percentage losing perimeter (%L) including the selected variables in the refined Random Forest analysis (Figure S6). Each cluster displays the average outcome. The value below the box indicates the percentage of data contained in the cluster. The bottom panel plots %L as a function of the 3D Perimeter and fractal surface dimension. The shades of blue and circle sizes are proportional to the level of %L. b) The two panels are analogous to a) but using the percentage of winning perimeter (%W) as an output variable. The percentage of winning is in this case proportional to the intensity of green.

For the percentage of winning outcome (%W), the regression tree selected the surface area (SA), 3D perimeter (P3D), and 3D perimeter to surface area ratio (P3D/SA) as the main nodes (Figure 4b). Corals with a large surface area, SA > 6482 cm^2^, had a higher %W and were classified on the right side of the tree. Among those, corals with small perimeter to surface area ratios, P3D/SA < 0.054 cm-1, formed the group with the largest percentage of winning perimeter, <%W> = 43%. On the left side of the tree, that is, SA< 6482 cm^2^, the secondary node was based on the 3D perimeter instead of the 3D perimeter to surface area ratio. Corals with large perimeters, P3D > 141 cm, formed the group with the lowest percentage of winning perimeter, <%W> = 14%. Figure S6b also plots the %W as a function of the surface area and the perimeter to surface area ratio. The sectors represent the regions selected by the regression tree. As expected, the bottom-right sector (large SA and small P3D/SA) had the highest value of percentage winning perimeter, while the top-left sector (small SA and large P3D) had the smallest percentage of winning perimeter. Notice that corals with larger %W resided in the bottom half of the scatter-plot, that is, the region with smaller perimeter to surface area ratio.

### Geometric predictors at the species level

The coral-algal competitive outcomes were also analyzed separately for species represented by more than five sampled colonies: *Orbicella faveolata* (n=12), *Montastraea cavernosa* (n=10), *Pseudodiploria strigosa* (n=8), *and Siderastrea sidereal* (n=7) (Figures S7, S8, S9, S10). For each species, the average percentage of losing perimeter (%L) as a function of the surface fractal dimension (D_s_) and 3D perimeter (P_3D_) (Figure S9) was compatible with the values obtained for the same regions in the global analysis (Figures 4a). This was also consistent for the percentage of winning perimeter (Figures S10 and 4b). Thus, the outcome averages for the regions selected in the global analysis led to equivalent results at the individual species level.

### Analysis of coral geometric properties across Curaçao regions

The corals sampled were grouped in three geographical regions: East, Central, and West (Figure S1). The four main geometrical indicators for the percentage of losing and winning interactions were compared using boxplots (Figure S11). The Central region showed the lowest value for the fractal dimension (median D_s_<2) and surface area (Figures S8a and S8b), indicating a higher percentage of corals losing against algae. The East region displayed a relatively large surface dimension, which was comparable to the West region (D_s_>2). Additionally, corals in the East region displayed the largest surface area of all.

## DISCUSSION

Coral geometrical properties are involved in the acquisition of resources as well as the defense of corals against benthic algae, and in this we were interested in determining if larger fractal dimensions and smaller perimeter-to-surface area ratios were favoring corals when facing competitive interactions with algae.

### Relation between coral geometry and coral-algal outcomes

Coral geometric properties explained 19–27% of the coral-algal interaction outcomes (Figure S6). The surface fractal dimension was instead the best single indicator for the percentage of perimeter that was losing or winning (p-value = 0.0021** and R^2^ = 0.18, Figure S5). This is consistent with the coral surface being essential for harvesting energy for growth and competition. To defend its perimeter, a coral colony depends on resources acquired through photosynthesis (carried out by endosymbiotic algae) and heterotrophic feeding (Porter, 1976). Losing corals had lower surface fractal dimensions (DS<2) and presented holes and large peninsulas, while winning corals had higher surface fractal dimensions (DS>2) and displayed more compact and rugose surfaces (Figure 3c). Higher perimeter-to-surface area ratios (P/SA) were correlated with winning corals as a secondary indicator when the surface area of corals was large enough (Figures 4b).

The multivariate statistical analysis selected the 3D Perimeter (P3D), fractal surface dimension (DS), and coral species as the most relevant variables for the percentage of losing perimeter (%L) (Figure S6a). These variables combined explained 19% of the variance of outcomes—similar to the variance explained by the surface fractal dimension alone, 18% (Figure S5a). For the percentage of winning perimeter (%W), the variables selected were the 3D perimeter to surface area ratio (P3D/SA), 3D perimeter (P3D), and surface area (SA) (Figure S6b). These variables combined explained 27% of the variance (Figure S5b). Low surface fractal dimensions, DS<2, were a good proxy for losing corals (Figure 4a), while large surfaces with low perimeter to surface ratios favored winning corals (Figure 4b).

### Implications of the fractal dimensions of coral colonies

Coral fractal dimensions have been used to differentiate coral species based on the structure and texture of corallites (Martin-Garin et al., 2007), characterize coral rugosity (Knudby and LeDrew, 2007), describe coral and sponge growth (Kaandorp & Kubler, 2001), measure coral mass over multiple scales (Basillais, 1997 & 1998), and distinguish functional groups such as coral rubble and algal flats on large reef scales (Purkis et al., 2005, 2006; Zawada & Brock, 2009). As shown in Figure 5, the perimeter of coral colonies (range 0.5 mm to 1 m) displayed fractal dimensions close to the topological dimension, D∼1 (current study). Larger colonies (range 0.1 m –100 m) had slightly larger values, D∼1.2 (Bradbury and Reichelt, 1983; Mark, 1984), and coral reefs (10 m – 5 km range) displayed values on the order of D∼1.5 (Purkis et al., 2006). The perimeters of seagrass beds and hard ground patches were similar, suggesting that the topography of the ground may be responsible for the increment of the fractal dimension. The surface fractal dimension of corallite sections adopted D∼0.8–1.0 at the septa range 0.1 mm – 1 mm (texture) and D∼1.2–1.6 at the calicular range 1 mm – 1 cm (structure) (Martin-Garin et al., 2007). The surface of coral colonies (1 mm – 1 m range) adopted fractal dimensions around the topological value, D∼1.85 – 2.15. Coral reefs (0.5 m – 5 km range) displayed larger values D∼2.28–2.61 (Zawada and Brock, 2009), which could be associated to the rugosity of the ground as in the case of the perimeter. Thus, the perimeter and surface fractal dimensions increase at larger scales.

**Figure 5.**
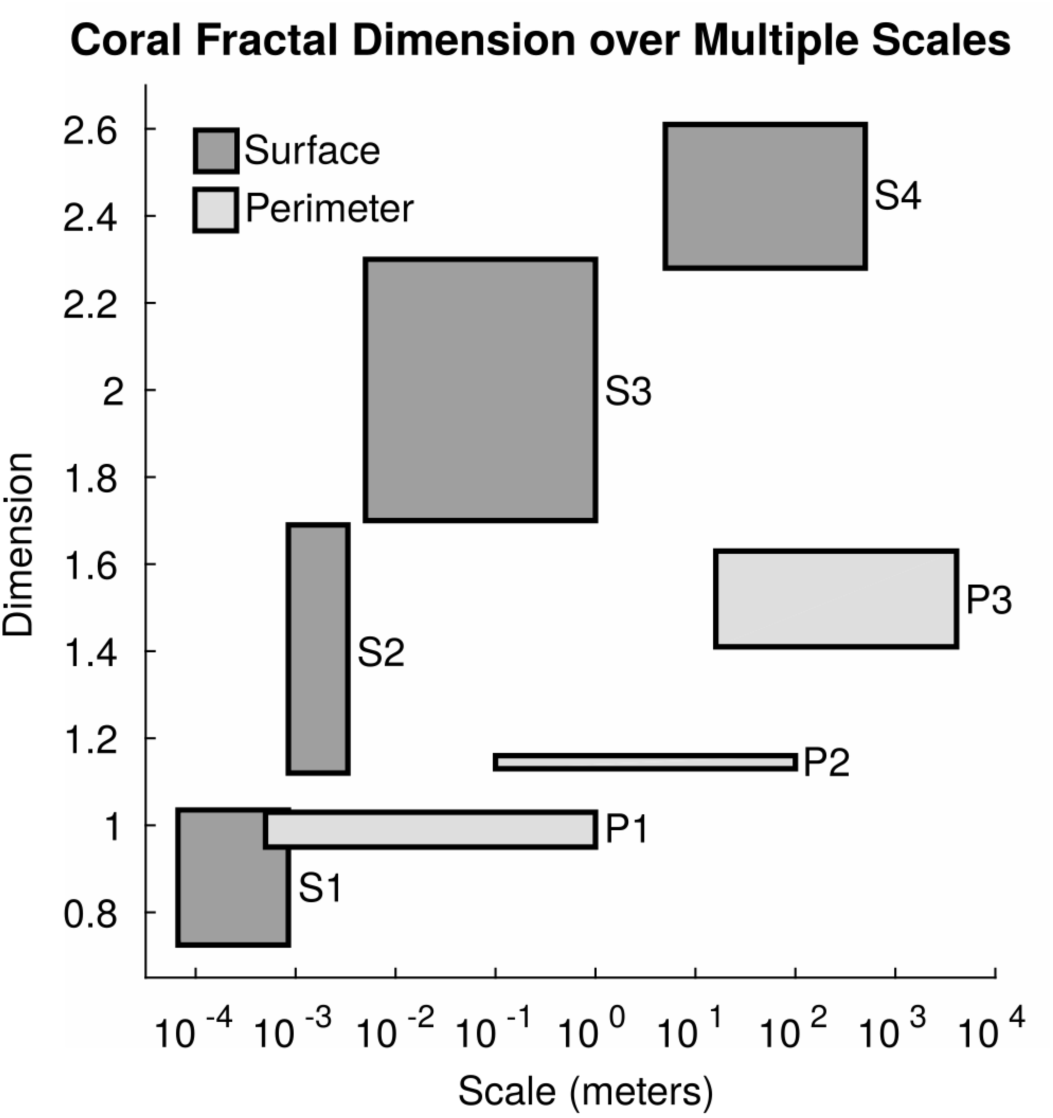
Coral fractal dimension over multiple scales. The chart plots the ranges of fractal dimensions measured across scales for different coral studies. The fractal dimensions are grouped in two categories: Surface fractal dimension (dark grey) and perimeter fractal dimension (light grey). For the perimeter, the ranges correspond to coral colonies (P1; current study), larger coral colonies (P2; Bradbury & Rachel, 1983; Mark, 1984), and coral reefs (P3; Purkis et al., 2006). For the surface, the ranges correspond to corallite texture (S1; Martin-Garin et al., 2007), corallite structure (S2; Martin-Garin et al., 2007), coral colonies (S3; current study), and coral reefs (S4; Zawada et al., 2009).

At the coral colony scale, the perimeter and surface dimensions were compatible with the Euclidean dimensions, D = 1 and D = 2, respectively (Figure 3). This justifies modeling coral colonies using Euclidean geometries (Meesters & Bak, 1996; Jackson, 1977; Naumann et al., 2009). The mean values of the surface fractal dimension, however, correlated with coral outcomes (Figure S5) and identified salient geometrical features. Corals with mean fractal dimensions smaller than two, DS < 2, displayed surfaces with holes and large peninsulas, while corals with fractal dimensions larger than two, DS > 2, displayed more compact surfaces with richer and more wrinkled textures (Figure 3c). Additional geometric metrics such as rugosity, vector dispersion, multivariate multiscale fractal dimension, and multifractal analysis (Reichert et al., 2017; Young et al., 2017, Chakraborty et al., 2016) might be necessary to refine the coral geometric analysis presented here.

The open regions observed in corals with a surface fractal dimensions smaller than two, D_s_<2, can represent more space for algae to occupy, thus leading to the DOC-Disease-Algae-Microbes (DDAM) positive feedback loop detrimental for those coral colonies (Dinsdale & Rohwer, 2011; Haas et al., 2011; Barott et al., 2012a, Roach et al., 2017). This lower fractal dimension associated to holes aligns also with the fact that corals have a limited capacity to regenerate lesions, and if they are larger than a certain size they may never be closed (Meesters et al., 1997). In fact, the sites sampled in the Central region of Curacao had a significantly lower surface fractal dimension than the East and West regions (see Figure S11). The combination of the geometrical properties and the decision trees (Figure 4) suggested that the East region is the healthiest region of Curaçao, followed by the West and Central regions. This analysis is consistent with field observations (Barrot et al. 2012c) and confirms the applicability of the geometrical analysis of corals as a proxy to assess coral-algal interactions.

## Conclusions and Perspectives

The geometrical properties of corals explained 19% to 27% of coral-algal competition outcomes. The perimeter and surface dimensions of coral colonies were non-fractal, but the mean surface fractal dimension displayed the strongest correlation with coral-algal interaction outcomes. Losing corals had low surface fractal dimensions (DS<2) and displayed holes and large peninsulas, while winning corals (DS>2) were more compact and displayed more rugose surfaces. Winning corals had larger surface areas with lower perimeter to surface area ratios, confirming that coral surfaces play a key energetic role in sustaining corals against algal attacks. The main geometrical predictors selected from the global analysis partitioned the percentage of losing and winning perimeters of individual species consistently. Additional data for individual species, however, will be necessary to confirm the relationship between geometrical properties and coral-algal interaction outcomes. Surveying the surface area and fractal dimensions of corals in other regions will help validate the generality of these results. Nevertheless, more sophisticated techniques such as multifractal analysis, might be necessary to understand why the surface fractal dimension is statistically non-fractal while displaying the strongest correlation discerning losing and winning corals. Additionally, it will be necessary to incorporate other descriptors that impact coral outcomes such as microbial and viral communities to achieve more accurate predictions.

## Acknowledgements

We acknowledge Mark Hate for the original artwork that was adapted to generate the method figure about the reconstruction of 2D and 3D corals. We also thank and acknowledge the support of our funding sources. The work of Forest Rohwer and Aaron Hartmann was funded by the PIRE grant: NSF Partnerships for International Research and Education Grant (1243541). Ty N.R. Roach work was supported by the National Science Foundation (G00009988).

